# RFcaller: a machine learning approach combined with read-level features to detect somatic mutations

**DOI:** 10.1101/2022.05.11.491496

**Authors:** Ander Díaz-Navarro, Pablo Bousquets-Muñoz, Ferran Nadeu, Sara López-Tamargo, Silvia Beà, Elias Campo, Xose S. Puente

## Abstract

**Motivation:** The cost reduction in sequencing and the extensive genomic characterization of a wide variety of cancers is expanding the use of tumor sequencing approaches to a wide number of research groups and to the clinical practice. Although specific pipelines have been generated for the identification of somatic mutations, their results usually differ considerably, and a common approach in many projects is to use several callers to achieve a more reliable set of mutations. This procedure is computationally very expensive and time-consuming, and it suffers from the same limitations in sensitivity and specificity as other approaches. Expert revision of mutant calls is therefore required to verify calls that might be used for clinical diagnosis. Machine learning techniques provide a useful approach to incorporate expert-reviewed information for the identification of somatic mutations.

**Results:** We have developed RFcaller, a pipeline based on machine learning algorithms, for the detection of somatic mutations in tumor-normal paired samples. RFcaller shows high accuracy for the detection of substitutions and indels from whole genome or exome data. It allows the detection of mutations in driver genes missed by other approaches, and has been validated by comparison to deep sequencing and Sanger sequencing. The pipeline is able to analyze a whole genome in a small period of time, and with a small computational footprint.

**Availability and implementation:** RFcaller is available at GitHub repository (https://github.com/xa-lab/RFcaller) and DockerHub (https://hub.docker.com/repository/docker/labxa/rfcaller).

**Supplementary information:** Supplementary data is available online.

## Introduction

During the last decade, the introduction of Next Generation Sequencing (NGS) has transformed the study of cancer, with the identification of hundreds of novel alterations driving tumor transformation^1^. Major international cancer projects such as the International Cancer Genome Consortium (ICGC)^2^ and The Cancer Genome Atlas (TCGA)^3^ have expanded the repertoire of genes mutated in cancer, as well as the biological processes involved in it^4–7^. The continuous reduction in sequencing costs, together with the clinical significance of certain mutations for prognosis or treatment decisions, has transformed the used of NGS from large sequencing consortia to small size laboratories and clinical centers. However, the utility of NGS relies on the availability of somatic mutation calling pipelines with enough sensitivity to detect most somatic mutations, and high specificity to prevent the calling of artifacts or germline variants as mutations.

Somatic single nucleotide variants (SSNVs) and small insertions/deletions (indels) constitute the most abundant type of mutation in tumor genomes, and different tools have been developed in order to call somatic mutations from tumor-normal paired samples. Most *state-of-the-art* variant callings are based in traditional statistical methods, such as CaVEMan^8^, MuTect2^9^, MuSE^10^, Strelka2^11^, Pindel^12^ or SMuFin^13^ among others. However, there is no consensus on the mutations detected by each caller, with a large number of private calls specific for each method. These differences are mainly due to the ability of each program to deal with the tumor heterogeneity and purity, normal contamination, sequencing and mapping artifacts, coverage, as well as different downstream filtering steps^14^. Due to the advantages of some pipelines to detect specific *bona fide* mutations, some collaborative projects such as the PanCancer Analysis of Whole Genomes (PCAWG)^5^ or the TCGA PanCancer Atlas MC3^15^, do not use a single caller but a combination of algorithms, keeping the intersection between them as the set of mutations that is more reliable. Despite the utility of this multi-pipeline approach to generate a consensus set of mutations, this strategy has a very large computational cost, demanding large servers and consuming up to days for the analysis of a single case.

In addition to classical statistical-based approaches, during the last years there has been an expansion in the use of machine learning strategies for different purposes^16,17^, including the development of new variant calling tools. The initial use of these methods was mainly focused on refinement, taking a list of potential variants extracted with other pipelines to filter and select a final set of mutations^18–20^. However, these approaches still have a negative influence on computing time and reproducibility. On the other hand, recently developed pipelines use other machine learning approaches^21,22^ or even neural networks^23,24^ to directly perform variant calling for somatic mutations, although in some cases the computing or installation requirements are too complex for a medium-sized laboratory or institution.

Here, we describe RFcaller, an accurate, fast, light computational requirements and easy-to-use tool that uses read-level features together with machine learning strategies to identify somatic mutations (SSNVs and indels) from normal-tumor paired samples. Our pipeline has been trained for whole genome sequencing (WGS) data and its results have been compared with those obtained by the PCAWG, being very similar to those resulting by combining several tools.

## Materials and methods

### Selection of somatic mutations

For the development of the algorithms, two different set of mutations were used, a training set and a testing set. To build them, we extracted with bcftools^25^ all possible somatic mutations from four WGS mantle cell lymphoma (MCL) samples sequenced at 30X coverage (M032 and M439 for training; M065 and M431 for testing). For the initial training, previously published mutations^26^ were defined as true positive mutations. With each iteration, all discordant calls were manually reviewed by three experts, through visual inspection, and the database was updated accordingly (Supplementary Table S1). This procedure resulted in the identification of novel *bona fide* mutations that would constitute false negatives in the initial set, as well as the rejection of certain mutations, such as artifacts or germline mutations present in the original dataset, that would represent false positives, respectively. After several rounds of training the algorithms and curating the set of mutations, all discordant variants had already been examined, which allowed us to obtain a reliable dataset for training and testing the final version of the algorithms.

### Algorithm training

To train the algorithms, we used the training set which contained 66,096 potential SSNVs (Supplementary Table S2) and 931 indels (Supplementary Table S3) for which read-level features were previously extracted (Supplementary Table S4). These data were used as input by TPOT^27^ (v0.11.1), with the default configuration of the *TPOTRegressor* function, to find the best pipeline to train the regression algorithms. As a result, an extremely randomized tree “Extra-Tree” Regressor for SSNVs and a Random Forest Regressor for indels were built. In both cases, a transformation of the data was carried out before the regression using the *StackingEstimator* function.

Once we had the algorithms, the test dataset, with 63,948 SSNVs and 2,506 indels (Supplementary Table S5), was used to select the best cutoffs for both pipelines. With this purpose, the result from RFcaller was filtered to get the “QUAL” field for those mutations that passed all filters (Supplementary Table S6). This parameter is calculated considering the initial quality from bcftools and the regression value for SSNV and indels, and only the regression value for homopolymer indels (polyindels):

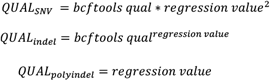

Then, ROC curves were generated and AUC metrics were calculated using the R package OptimalCutpoints^28^ with the *MaxEfficiency* method. False/True Positive/Negative ratios were calculated using the formulas described in the ROCR^29^ R package.

### Computational cost

To compare the performance of RFcaller with other *state-of-the-art* tools, the docker container corresponding to the four callers used by PCAWG for the detection of SSNVs was downloaded (https://dockstore.org/organizations/PCAWG/collections/PCAWG). After minor fixes of broken links in the Sanger and DKFZ tools, all of them were run with the default parameters for one random donor. In case the tools allowed to choose the number of threads and RAM to be used, 20 threads and 200 Gb of memory were specified. In addition, because RFcaller allows multiple samples to be run simultaneously, four cases were run in parallel using the default parameters to calculate the computational cost. To improve data interpretation, some axes were broken using the R package ggbreak^30^.

### PCAWG analysis

To validate that the trained models are applicable for liquid and solid tumors and to compare the results to those obtained by the PCAWG pipeline, RFcaller was run for the CLLE-ES and BRCA-EU studies, with an average tumor coverage of 30X and 50X, respectively, and 30X for normal samples in both. PCAWG BAM files were downloaded from the “collaboratory” repository using the score-client program (Supplementary Table S7). RFcaller was run with its default parameters for all samples and the obtained results were combined into a single VCF file for each study. A custom panel of normals was used to annotate variants in complex regions. The set of mutations detected by the PCAWG pipeline were extracted from the controlled consensus callsets for SSNV/Indel. To analyze coding and non-coding mutations, the Variant Effect Predictor (VEP) tool^31^ was launched for both datasets using the following options: `--offline --format vcf --dir_cache homo_sapiens —symbol --force_overwrite --total_length --numbers --ccds --canonical --biotype --pick --vcf --assembly GRCh37`.

To be able to compare both set of mutations in the most accurate manner: (i) dinucleotides and trinucleotides from RFcaller were split as this feature is not available for PCAWG, (ii) RFcaller mutations located in alternative chromosomes and PCAWG’s variants that appear in our custom dbSNP were removed and (iii) only mutations that passed all filters were studied. For this comparison, a mutation was considered as subclonal when its variant allele frequency (VAF) was lower than 0.15, in accordance with the sensitivity of Sanger sequencing.

For the purpose of calculating the precision and recall for both pipelines in each study, 1% or at least 50 discordant mutations from each section were manually reviewed by a panel of experts. Thus, a total of five blocks were checked: mutations detected only by RFcaller and mutations detected between one and four of the callers used by the PCAWG, as the ratio of false positives may be different between them. The results obtained were then extrapolated to the whole set of mutations in order to calculate the parameters needed to define precision and recall for both pipelines. These measures were calculated with the *prediction* and *performance* functions of the R package ROCR^29^.

Additionally, two bases upstream and downstream of the mutations were selected to reconstruct the context and generate a matrix with which extract the mutational signature of RFcaller and PCAWG-private mutations and those detected by both pipelines. The sigminer^32^ R wrapper (nrun = 300 and refit = TRUE) was selected to run the SigProfilerExtractor framework^33^.

Finally, deep sequencing data generated by previous studies^34,35^ for some CLLE-ES cases (Supplementary Table S8) were used to analyze possible subclonal mutations in driver genes. In order to compare both results, only mutations in CLL driver genes and donors analyzed by both WGS and deep sequencing were selected. In addition, mutations detected by deep sequencing were removed from the analysis if they were germline or there was insufficient coverage or reads supporting the mutation by WGS (Supplementary Table S9).

### Sanger validation

To perform verification of private calls obtained from the analysis of CLLE-ES cases, five and two mutations detected only by RFcaller and PCAWG, respectively, were chosen to be verified by Sanger sequencing. These positions were chosen because they appeared in known driver genes for CLL and because tumor and/or normal DNA was available. The list of primers and melting temperatures are listed in Supplementary Table S10.

### Exome analysis

To test the performance of RFcaller on exome sequencing data, we selected five CLLE-ES cases previously analyzed by WGS and for which exome data were available (Supplementary Table S8). RFcaller was run with default parameters and LIKELY_GERMINAL variants were removed. Only mutations within the targeted regions of the exome (Agilent – SureSelect Human All Exons V4) were taken into account. Finally, for those mutations not detected by both methods, total coverage and number of mutated reads were extracted in order to determine the cause for loss.

## Results

### RFcaller workflow

An overview of the RFcaller’s workflow is provided in Figure 1. The pipeline takes as input the BAM files from the normal-tumor paired samples and starts performing a basic variant calling using bcftools (v1.10.2) with the -P option set to 0.1 to enable calling of low frequency variants. Then, indels are normalized, and common SNPs (dbSNP v153), and variants within five base pairs of an indel, are removed. To increase the speed of the pipeline, low quality calls are filtered (<15 for SSNVs and <40 for indels). Remaining mutations are divided into three different files to be processed independently: SSNVs, short-indels (<7bp) and long-indels (≥7bp).

**Figure 1.**
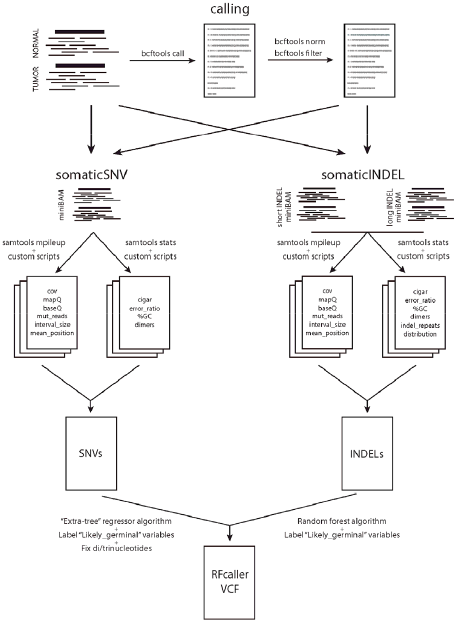
Flowchart of the RFcaller pipeline.

SSNVs and indels have a specific pipeline where read-level features are extracted for those mutations that meet basic requirements that can be customized, such as minimum coverage (≥7), maximum number of mutated reads in normal (≤3 for SSNVs and ≤2 for indels) or a minimum number of mutant reads in tumor (≥3 for SSNVs and ≥4 for indels). These filters were chosen because positions that fail to meet these requirements cannot be confidently classified as *bona fide* mutations from the available data. Once all features have been extracted, a CSV file is generated to be used by the algorithm. The result is a VCF file with mutations that have passed the threshold for the “QUAL” field.

To classify mutations that might be germinal but have passed the previous filters, a 95% confidence interval is applied to calculate the expected number of mutant reads in normal, considering: the VAF of the mutation in tumor sample, the expected contamination of tumor in normal sample defined by the user and the normal coverage. Thus, if the number of mutated reads in normal is greater than the expected, the position is labeled as “LIKELY_GERMINAL”.

Finally, the RFcaller pipeline for SSNVs searches for dinucleotides or trinucleotides mutations within the results. With this step, if two mutations are found together in the same allele, they are merged into a single mutation to be accurate when predicting its functional effected, a step that is usually missed by most commonly used somatic callers.

### RFcaller training

For the initial training step, previous results from the genomic analysis of two mantle cell lymphomas^26^ were used to annotate the set of mutations, and RFcaller was trained with this initial dataset. The obtained results were compared with those used for training, and all discordant positions were manually reviewed to improve the accuracy of the dataset. These steps were repeated until all discrepancies were classified by an expert panel. After that, 2,208 and 2,901 calls were reviewed for training and testing, respectively, resulting in a high quality set of mutations to train and test the final versions of the algorithm (Table1).

**Table 1.** Number of total and manually reviewed mutations used for training and testing RFcaller. TP: Number of True Positive mutations; TN: Number of True Negative mutations

|  | Training set |  |  |  | Test set |  |  |  |
| --- | --- | --- | --- | --- | --- | --- | --- | --- |
|  | SSNV |  | Indel |  | SSNV |  | Indel |  |
|  | TP | TN | TP | TN | TP | TN | TP | TN |
| Manually reviewed | 915 | 730 | 321 | 242 | 924 | 959 | 528 | 490 |
| Total | 8,362 | 57,734 | 504 | 427 | 6,909 | 57,039 | 696 | 1,810 |

In order to select the best cutoff for the pipeline, SSNVs, indels and homopolymer indels were considered independently as they represent mutations whose detection is influenced by different features. The separation between both types of indels (isolated or within a homopolymer trait) was introduced due to the bias of the initial calling performed by bcftools against indels within homopolymeric tracts, giving very low scores to mutations that otherwise appear to be real. Furthermore, different formulas were considered to calculate the “QUAL” threshold used by RFcaller (Supplementary Table S11).

Although the RFcaller score provided high accuracy, we observed that by combining the regression obtained by RFcaller with the score given by bcftools, the accuracy was improved over each one independently, suggesting both scores complement each other. We did not observe major differences between formulas for SSNVs and indels according to the area under the curve metric (AUC), so we selected the formulas with the highest F1 score. Thus, the cutoffs were 10.726 for SSNVs, 32.1418 for indels and 0.7723 for homopolymer indels (Supplementary Figure S1), which achieved 1.3%, 7.18% and 8% of false positive mutations, respectively. We observed that many of the false positives belonged to complex regions like microsatellites or GC-rich sites, appearing also in normal samples from other donors. Therefore, we used a panel of normals to filter these calls and improve the accuracy of the pipeline.

In terms of the number of variables selected, only 16 and 27 read-level features were considered for SSNVs and indels respectively, which helped us to avoid overlapping features that can be counterproductive and lead to overfitting. Another important aspect we considered during the selection of these features was the difficulty by which they can be extracted, resulting in a fast pipeline for medium-size servers. Thus, the analysis of four WGS tumor-normal paired samples using 20 threads consumes only ~5 GiB of RAM and takes ~3 hours/case, while using only 10 processors the analysis is extended up to ~4.5 hours/case (Supplementary Figure S2d).

When RFcaller was compared with the callers used by PCAWG for the detection of SSNVs, only the muse-variant-caller (~2.5 h) was faster than RFcaller (~4.8 h) (Supplementary Figure S2), while sanger-variant-caller was the slowest, taking more than 70 h for a single case. In terms of memory consumption (RSS), mutect-variant-caller is the most demanding, consuming between 100 GiB and 250 GiB during half the time it is running (~5 h). In this case, RFcaller and muse-variant-caller consume the least memory with an average of 5 GiB. It is important to note that although we have used the SSNVs specific callers, all of them, except MuSE, also detect indels, which would imply that RFcaller is the fastest and least resource consuming tool for the simultaneous calling of SSNVs and indels.

### Validation of RFcaller pipeline: PCAWG analysis

To test RFcaller against a validated set of cancer WGS cases we used data from the PCAWG study belonging to two different projects (CLLE-ES and BRCA-EU), representative of liquid and solid tumors, with a total of 89 and 75 cases, respectively (Supplementary Table S8). RFcaller results were compared to those mutations labeled as “PASS” by the PCAWG mutation calling pipeline. Due to the inherent differences between SSNVs and indels, we performed each analysis independently.

#### Somatic Single Nucleotide Variants

After merging RFcaller and PCAWG “PASS” mutations, we observed that ~70% of SSNVs were detected by both pipelines in both studies. However, and even though the number of shared mutations was almost the same, for samples from the CLLE-ES project 11% of mutations were detected only by the PCAWG pipeline vs. 16.3% mutations specifically detected by RFcaller. For BRCA-EU-derived mutations, only 4.4% mutations were RFcaller-specific, vs. 25.4% for PCAWG pipeline (Figure 2a). A detailed analysis of those differentially called mutations revealed that the mean VAF for SSNVs detected by both pipelines was 0.41 and 0.27 for CLLE-ES and BRCA-EU, respectively (Figure 2b). However, those detected by the PCAWG pipeline but not RFcaller had a mean VAF of 0.16 and 0.10 for CLLE-ES and BRCA-EU, respectively (Figure 2b), suggesting that they constitute subclonal mutations. In fact, only 29% and 50% of them could be detected by more than two callers in the PCAWG pipeline for CLLE-ES and BRCA-EU, respectively (Figure 2c). Furthermore, those SSNVs detected by RFcaller but not the PCAWG pipeline had a mean VAF of 0.46 for CLLE-ES and 0.28 for BRCA-EU, similar to those detected by both pipelines, suggesting that they constitute clonal mutations detected by RFcaller. Some of them showed minor tumor in normal contamination (1-3 mutant reads), common in hematological tumors, resulting in most callers missing these true positive somatic mutations, while RFcaller is able to retain most of them.

**Figure 2.**
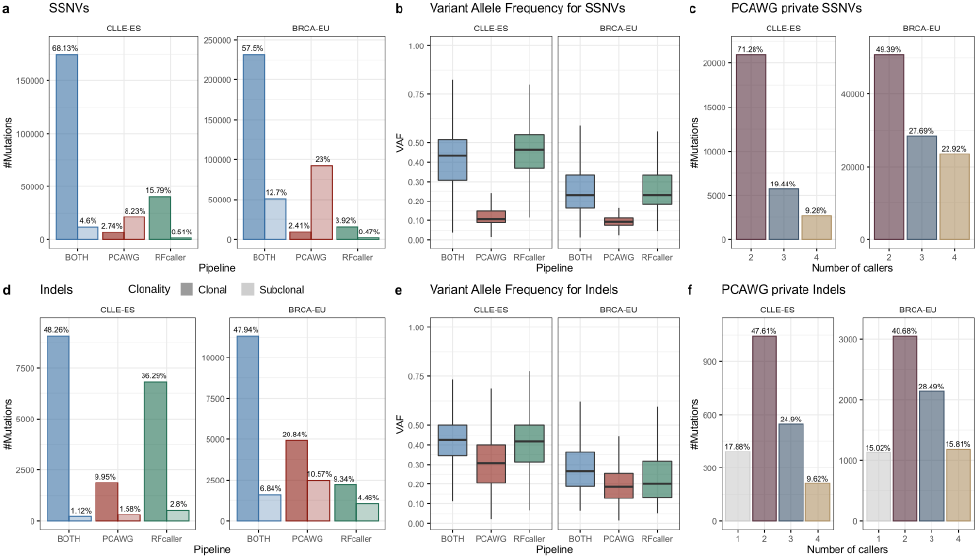
Summary of mutations detected by PCAWG and/or RFcaller pipelines for SSNVs and indels. a,d) Classification of mutations according to the pipeline that can detect them. Mutations are divided in clonal (VAF≥0.15) and subclonal (VAF<0.15) mutations. b, e) Distribution of the variant allele frequency of the mutations identified by both pipelines, or specifically by RFcaller or PCAWG pipeline. c, f) Number of callers detecting each of the PCAWG-private mutations.

To explore the set of discordant mutations between both pipelines, we randomly selected 1-2% of the pipeline-private calls (n=776 for CLLE-ES and n=1233 for BRCA-EU) to be manually reviewed by a panel of experts (Supplementary Table S12 and Supplementary Figure S3). As expected, PCAWG-specific variants detected by four callers are more precise than those identified by two tools (Table 2). Surprisingly, the difference in precision for RFcaller-private mutations between studies was very high, 98.5% for CLLE-ES and 74.5% for BRCA-EU, probably reflecting the fact that RFcaller was trained using a hematological tumor. However, despite the apparently higher number of false positives, RFcaller-private calls only represent 18.3% and 5.9% of the total number of SSNVs detected by RFcaller in the CLLE-ES and BRCA-EU projects, respectively. Considering the observed number of false positive calls within these sets, the real precision of RFcaller calls for SSNVs is 99.7% and 98.5% for CLLE-ES and BRCA-EU, respectively, while the precision of the PCAWG pipeline is 97.3% for both studies (Figure 3).

**Figure 3.**
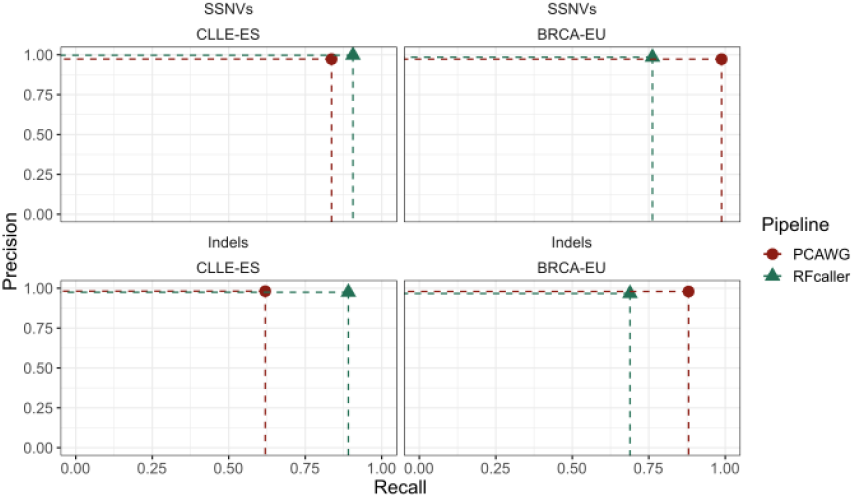
Accuracy of RFcaller and PCAWG pipelines for SSNVs and indels against CLLE-ES and BRCA-EU datasets. RFcaller shows a higher recall in both SSNVs and indels for CLLE-ES, whereas in BRCA-EU the PCAWG manages to detect a higher number of mutations. The precision of the two pipelines is similar in all conditions.

**Table 2.**
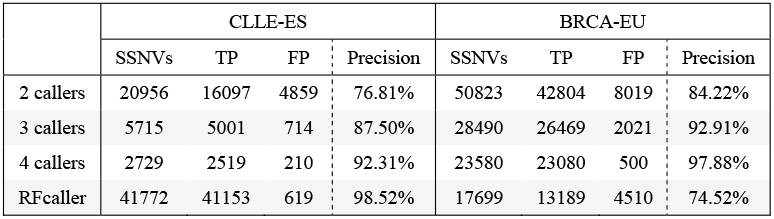
Total number of false positive private SSNVs extrapolated after manual revision. TP: Number of True Positive SSNVs; FP: Number of False Positive SSNVs

To further explore these private mutations, we extracted the mutational signatures independently for the set of mutations detected by both pipelines, as well as for those specific for each caller (Supplementary Figure S4). We could see that in CLLE-ES study, both RFcaller-private SSNVs and those common to both pipelines contained the same signatures (SBS1, 5, 8 and 9), while PCAWG-private SSNVs shared 3 signatures (SBS1, 5 and 8), missed one (SBS9) and contained two signatures not detected in the common set (SBS23 and 51), although affecting a limited number of samples. While on the BRCA-EU study, both pipelines missed some signatures present in the common set (2 PCAWG and 4 RFcaller), and both detected one and two signatures, respectively, not present in the common. Together, these results suggest that the private mutations detected by RFcaller constitute *bona fide* calls, with a similar profile to those detected by both pipelines.

#### Small insertions/deletions

The analysis of small indels revealed that there were more differences between pipelines than those seen for SSNVs. In this regard, only ~50% of indels were detected by both RFcaller and PCAWG pipelines, however for CLLE-ES RFcaller-private calls represented 39.1% of the total number of indels whereas only 11.5% of them were PCAWG-specific. In contrast, in BRCA-EU, RFcaller and PCAWG-private mutations accounted for 13.8% and 31.4% respectively (Figure 2d). Moreover, among them, less than 45% of PCAWG-private indels were detected by more than two callers (Figure 2f), reflecting the difficulty to identify somatic indels in tumor samples.

To further explore pipeline-private indels, we selected at least 50 indels from each group for expert review (n=283 for CLLE-ES and n=429 for BRCA-EU) (Supplementary Table S12 and Supplementary Figure S3). We observed that the precision within PCAWG-private indels was very high, varying between 70% and 99% depending on the number of individual callers supporting the call (Table 3). In contrast, the precision observed for RFcaller was 89%, despite the fact that the total number of indels detected by this pipeline was much higher. Similar to SSNVs, the observed VAF was slightly higher in CLLE-ES compared to BRCA-EU (0.42 vs 0.29), probably reflecting higher tumor purity. Nonetheless, we did not observed differences in VAF between pipeline-private indels (Figure 2e), suggesting that pipeline-specific mutations were not due to clonality, as they were for SSNVs, but to other factors such as alignment issues, size of the indel, the presence of microsatellites or if they were within homopolymer tracks. Despite the higher precision obtained by the PCAWG pipeline for indel calling, this might be at the expense of a larger number of false negative calls in otherwise clonal and *bona fide* somatic indels, as shown by the number of true positive calls detected by RFcaller (Figure 3).

**Table 3.** Total number of false positive indels extrapolated after manual revision. TP: Number of True Positive indels; FP: Number of False Positive indels

|  | CLLE-ES |  |  |  | BRCA-EU |  |  |  |
| --- | --- | --- | --- | --- | --- | --- | --- | --- |
|  | Indels | TP | FP | Precision | Indels | TP | FP | Precision |
| 1 caller | 392 | 276 | 116 | 70.37% | 1125 | 1008 | 117 | 89.57% |
| 2 callers | 1044 | 1002 | 42 | 96.00% | 3046 | 2792 | 254 | 91.67% |
| 3 callers | 546 | 495 | 51 | 90.74% | 2133 | 2097 | 36 | 98.31% |
| 4 callers | 211 | 207 | 4 | 98.18% | 1184 | 1174 | 10 | 99.14% |
| Rfcaller | 7335 | 6916 | 419 | 94.29% | 3257 | 2721 | 536 | 83.54% |

In the case of mutational signatures detected for indels (Supplementary Figure S4), private calls from both pipelines contained most of the mutational signatures present in common mutations, while in the case of RFcaller-private indels, 3 signatures not present in the common set were detected for both CLLE-ES and BRCA-EU studies.

### Exome analysis

RFcaller was trained with WGS data, but as the features used for the prediction are at read level, this pipeline could also be used for exome analysis. In order to test the ability of RFcaller to detect mutations by WES, exomes from five cases previously analyzed by WGS were run with default parameters. Results were compared with those obtained by RFcaller and PCAWG in the WGS analysis after filtering for mutations within target regions in WES. Thus, 63% (n=110) of mutations detected by WES were also detected by WGS. Additionally, we were able to identify 47 novel mutations for which there was neither coverage nor any mutated read in WGS (Figure 4a). When we made the comparison in the opposite direction, we found that 55% (n=136) of the mutations detected by WGS did not appear by WES. However, 93% (n=126) of these missing mutations had no coverage or any mutated read in the exome or were clearly germinal (Figure 4b). Only 10 mutations detected by WGS had enough coverage in WES and were not detected, constituting false negatives (RFcaller exome recall = 94%). Similarly, considering the 17 mutations that were labeled as germinal by WGS but detected by WES as false positives, RFcaller achieves a precision of 90% (Supplementary Files).

**Figure 4.**
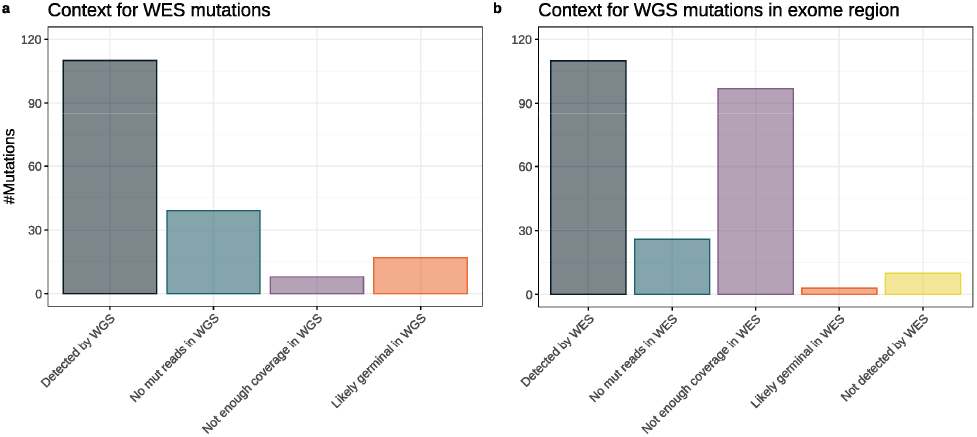
Comparison of mutations detected by analysis of WGS and WES in selected donors. Comparison is limited to exomic regions. a) Mutations detected by WES and analysis of their status in WGS. b) Mutations detected by WGS and analysis of their status in WES samples.

### Detection and verification of mutations in driver genes

From the above data we can conclude that RFcaller has a similar accuracy to detect SSNVs, and an increased sensitivity to detect indels at the cost of a smaller specificity. To explore if these differences might allow the detection of previously missed mutations with potential clinical impact, we analyzed somatic mutations on the set of driver genes previously described in these two tumor types (Knisbacher, B et al. 2022 (in press))^36^ (Supplementary Table S13). This analysis resulted in the identification of 155 coding mutations in driver genes in the CLLE-ES project and 162 in the BRCA-EU study. Out of those calls, 83% of them were shared between both pipelines, while 53 (17%) in 35 driver genes, were pipeline-specific (Supplementary Table S14).

Those pipeline-specific mutations were manually reviewed, resulting in the identification of 19 clonal mutations detected by RFcaller (12 SSNVs and 7 indels) vs. 4 clonal SSNVs detected by the PCAWG pipeline in CLLE-ES. For the BRCA-EU project, 8 clonal mutations were detected by RFcaller (5 SSNVs and 3 indels) vs. 4 clonal detected by PCAWG pipeline (3 SSNVs and 1 indel).

For seven private calls detected in the CLLE-ES study (5 by RFcaller, and 2 by the PCAWG pipeline), tumor and normal DNA was available for verification by Sanger sequencing, except two cases in which only tumor DNA was available (Supplementary Table S8). This analysis resulted in the verification of all RFcaller-private calls (Supplementary Figure S5), as well as one of the PCAWG-private SSNVs. The last call could not be verified because it was a subclonal mutation with a very low VAF (8.7%), that falls below the detection limit of this technique.

To further perform an orthogonal validation of these pipelines, we took advantage of a previous study in which 26 CLL driver genes had been analyzed by deep-sequencing in some of the CLL cases used by PCAWG^34,35^. A total of 77 mutations, excluding germline calls, were detected in 28 cases, for which enough coverage was available in WGS to make a call (Supplementary Table S9). Due to the high depth of sequencing, VAF was very variable (range 0.0029 to 0.9665), therefore, mutations were classified as clonal if VAF≥0.15 (n=44, median 0.43), and subclonal if VAF<0.15 (n=33, median 0.03). As expected, most subclonal mutations could not be detected from WGS data, as each pipeline was only able to detect 6/33 subclonal mutations (18%). By contrast, most clonal mutations detected by deep sequencing could be also identified by RFcaller (39/44, 89%), while the performance of the PCAWG pipeline was slightly lower (31/44, 70%). The mutations specifically detected by RFcaller affected *NOTCH1* (3), *ATM* (2), *TP53, RPS15, MGA* and *DDX3X*, some of which have been associated with poor prognosis and whose presence might impact clinical decisions. The PCAWG pipeline was able to identify a mutation in *ATM* that was not detected by RFcaller due to a very low VAF (0.065). Together, these results support the utility of RFcaller to identify novel clonal driver mutations of potential clinical value.

## Discussion

The application of NGS techniques for clinical diagnosis in tumor samples requires procedures that provide enough sensitivity and specificity, while at the same time do not require large computing resources to achieve the analysis in a reasonable amount of time. To increase accuracy, a final step of manual review through visual inspection is usually carried out for mutations that might be clinical informative. This manual revision increases the specificity, but at the cost of a labor intensive process. Recent advances in machine learning approaches are suitable to incorporate features that experts consider when distinguishing between *bona fide* mutations and false positives. However, most available programs that use machine learning approaches for somatic mutation calling have been trained with high depth of coverage WES using *in silico*^20,21^ or orthogonal validated mutations^22^ and cannot be used for whole genome analysis.

In this work, we have taken advantage of a manually curated dataset of real mutations with features that an expert curator might consider when manually reviewing a mutation in a research or clinical context. Thus, we have achieved a very high sensitivity to detect SSNVs and small indels, while at the same time maintaining a low footprint, with low CPU and RAM consumption, being able to analyze a whole genome in less than 5 hours. Moreover, although it has been trained with WGS data, it has shown a good performance in exome samples.

On the other hand, even though our selected features are often used by similar programs, they process SSNVs and indels in the same manner, when clearly the two types of mutations have different characteristics. In this regard, we analyzed SSNVs and indels separately, which allowed us to detect indels with higher accuracy without affecting the ability to detect SSNVs. Indeed, we have shown that RFcaller performance is similar to that of a combination of complex pipelines used in the PCAWG project to detect clonal mutations, with the ability to detect new ones, some of them in driver genes, what might contribute to improve the detection of actionable mutations^37^. Furthermore, we showed that RFcaller is able to detect mutations even in the presence of some tumor contamination in the normal sample, a common problem in some hematological tumors that might lead to false negatives with other pipelines. Finally, we have demonstrated that most RFcaller false negatives were subclonal mutations with very low VAF, whose analysis might require additional tools and might not be as critical for taking clinical decision.

In conclusion, we have developed a pipeline called RFcaller, that is provided under a Docker system, which allows its easy and fast installation without version incompatibilities. This tool allows the identification of clonal mutations with the same efficiency as *state-of-the-art* pipelines, but with a smaller footprint in computing resources.

## Data availability

RFcaller and the scripts used to train the algorithms are available at the GitHub repository (https://github.com/xa-lab/RFcaller), and a docker with all the requirements and necessary files to run the pipeline has been built to improve reproducibility and facilitate the use of the program (https://hub.docker.com/repository/docker/labxa/rfcaller). Additionally, the scripts with the files we have used to obtain the results shown above can be found in the supplementary material.

## Funding

This work was supported by Ministerio de Ciencia e Innovación [SAF2017-87811-R to XS.P., PID2020-117185RB-I00 to XS.P.]; Fundación Asociación Española Contra el Cáncer (AECC) [to XS.P. and S.B.]; Centro de Investigación Biomédica en Red Cáncer (CIBERONC) [to XS.P. and E.C.]; Instituto de Salud Carlos III and co-funded by European Union (ERDF/ESF, “Investing in your future”) [PMP15/00007 to E.C., PI17/01061 to S.B.]; “La Caixa” Foundation CLLEvolution [HR17-00221 to E.C.]; Ministerio de Economía y Competitividad (MINECO) [RTI2018-094274-B-I00 to E.C.]; Generalitat de Catalunya AGAUR [2017-SGR-709 to S.B., 2017-SGR-1142 to E.C.]; Department of Education of the Basque Government [PRE_2017_1_0100 to A.D-N.]; Asturian Government [to S.L-T.]; 2021 AACR-Amgen Fellowship in Clinical/Translational Cancer Research [21-40-11-NADE to F.N.], the European Hematology Association (EHA) Junior Research Grant 2021 [RG-202012-00245 to F.N.], and Lady Tata Memorial Trust (International Award for Research in Leukaemia 2021-2022) [LADY_TATA_21_3223 to F.N.]. E.C. is an Academia Researcher of the “Institució Catalana de Recerca i Estudis Avançats” (ICREA) of the Generalitat de Catalunya. IUOPA is funded by the Asturian Government and Fundación Cajastur.

### Conflict of Interest

X.SP. is co-founder and equity holder of DREAMgenics. E.C. has received research funding from Gilead Sciences, and has been consultant for Takeda, Celgene and Gilead, and is author in a Lymphoma and Leukemia Molecular Profiling Project (LLMPP) patent “Method for selecting and treating lymphoma types” PCT/US14/64161.

## Author contributions

A.D-N. developed the software, performed the bioinformatical work, interpretated data, designed the figures and wrote the manuscript. P.B-M., F.N. and S.L-T. contributed to data analysis and performed experimental work. S.B. and E.C. collected samples and interpreted data. X.S.P. designed the study, interpreted data, wrote the manuscript, and directed the research. All authors read, commented on, and approved the manuscript.

## Supplementary Figure Legends

**Supplementary Figure S1.**
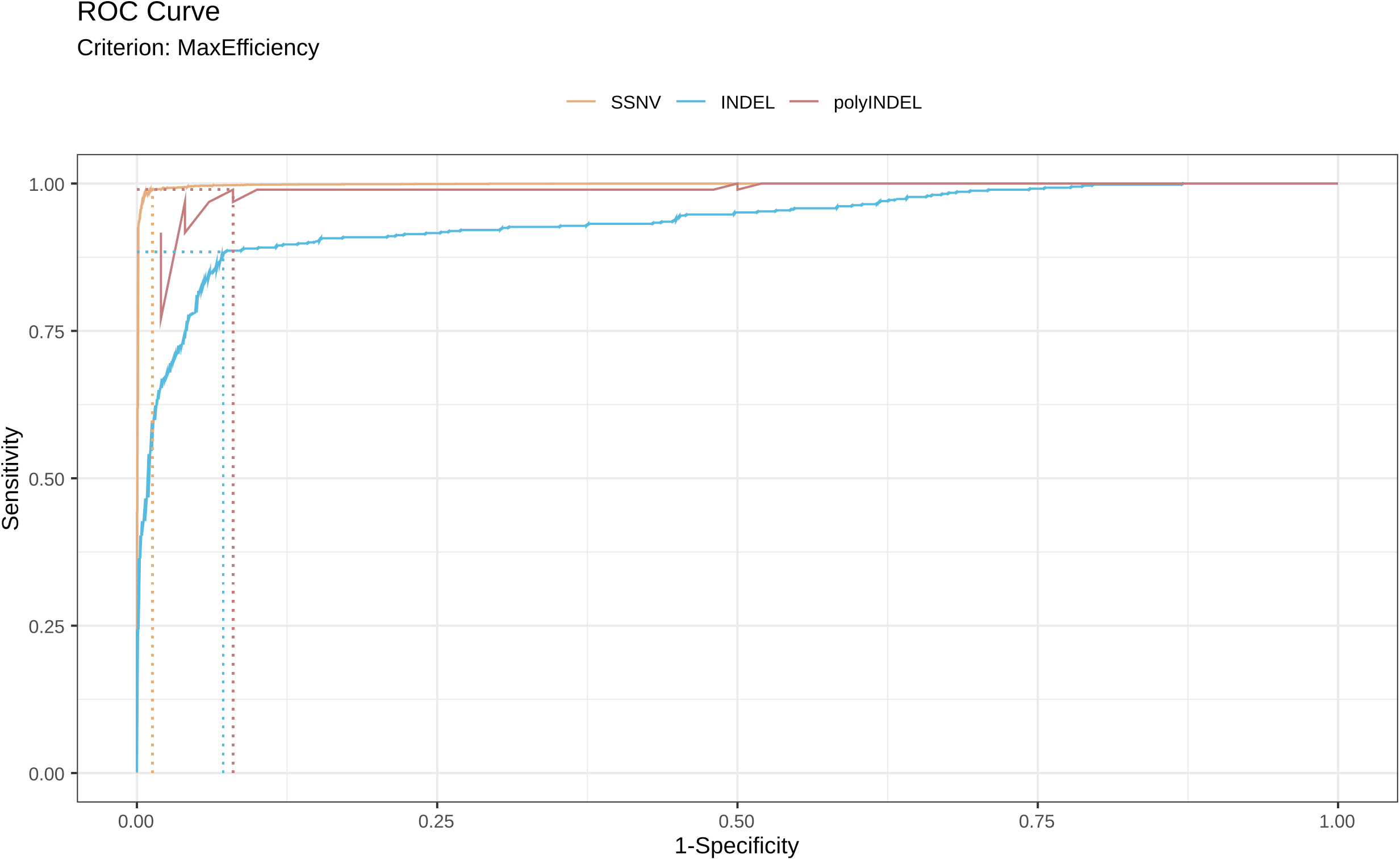
ROC curves for each mutation type with RFcaller results for the test data set using the formulas with the best F1 score. Cutoffs were obtained with the MaxEfficiency criterion.

**Supplementary Figure S2.**
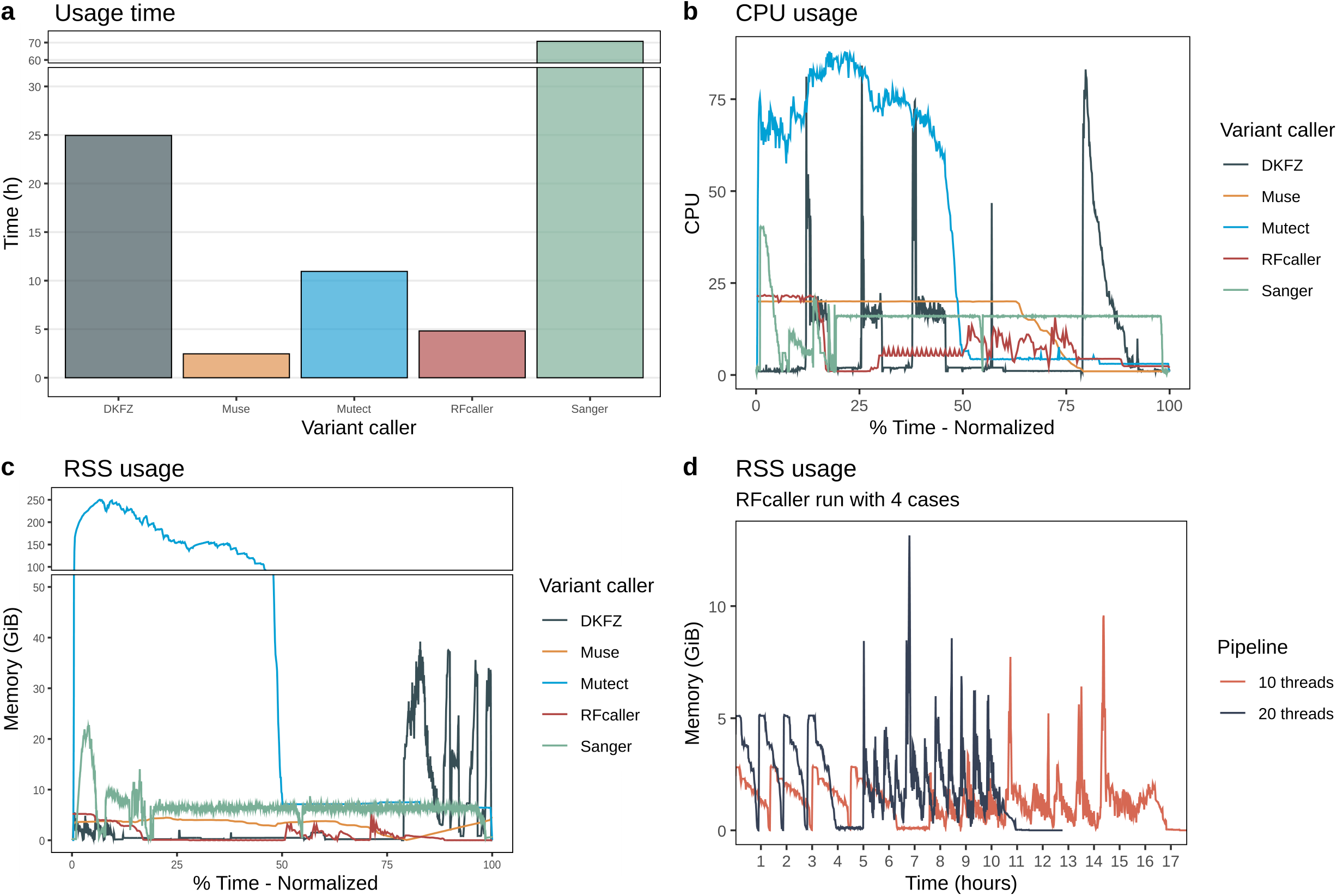
Representation of a) the time it takes for each caller to complete a case. b, c) CPU and memory (RSS) usage for the duration of the pipelines (RSS memory was limited to 200Gb when the caller allowed it). d) memory (RSS) usage of RFcaller pipeline when it is run with 10 or 20 threads for four independent cases simultaneously.

**Supplementary Figure S3.**
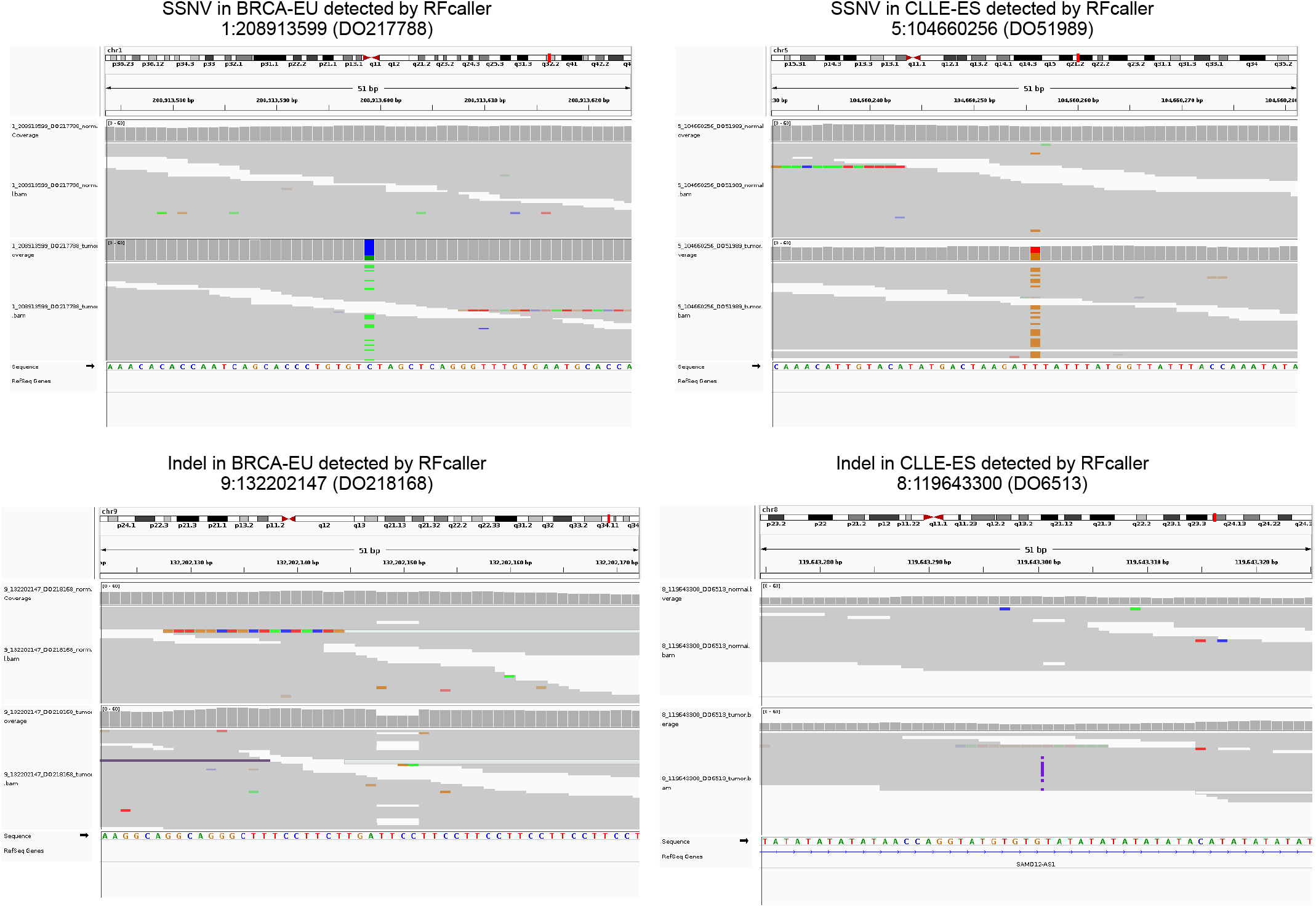
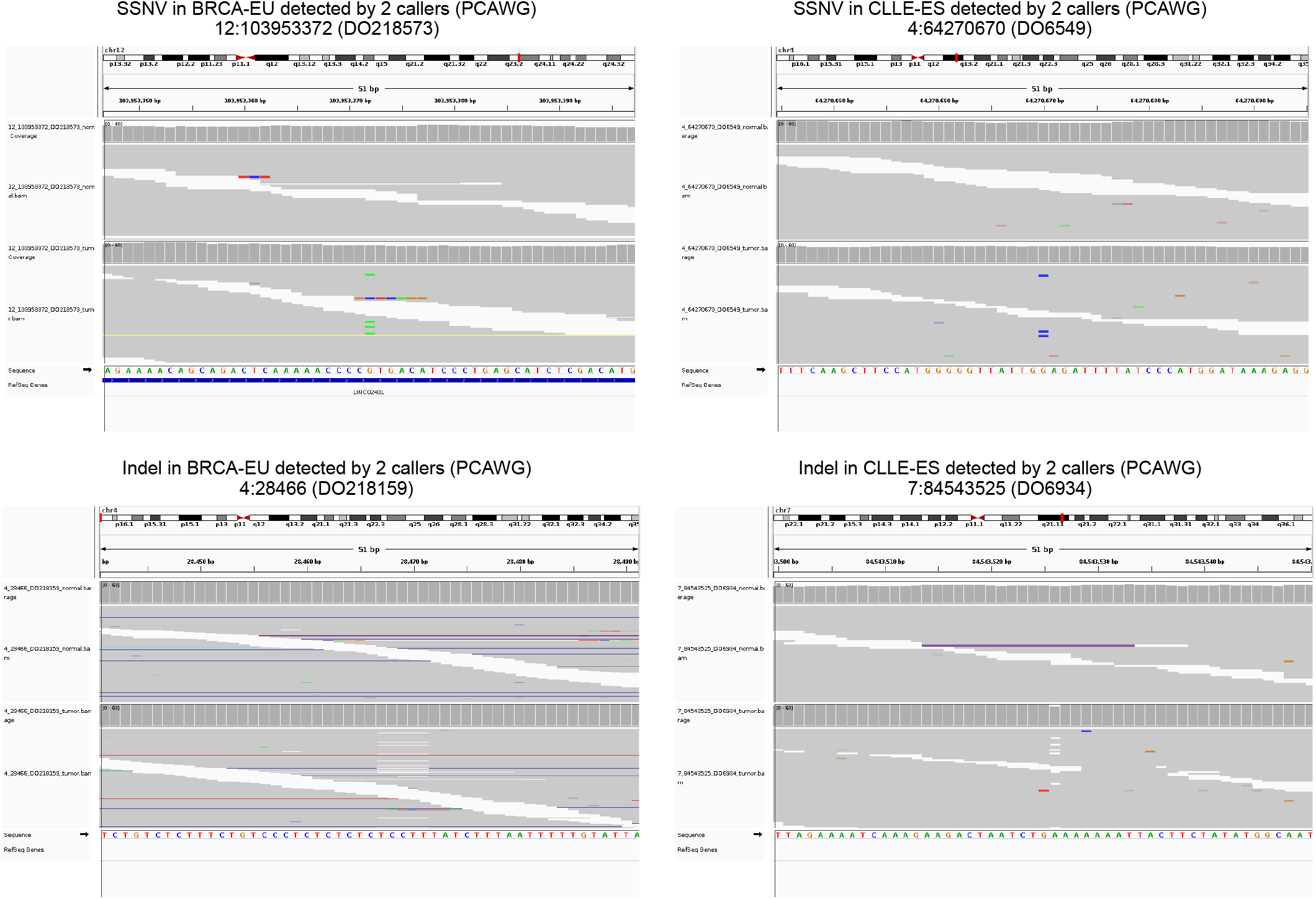
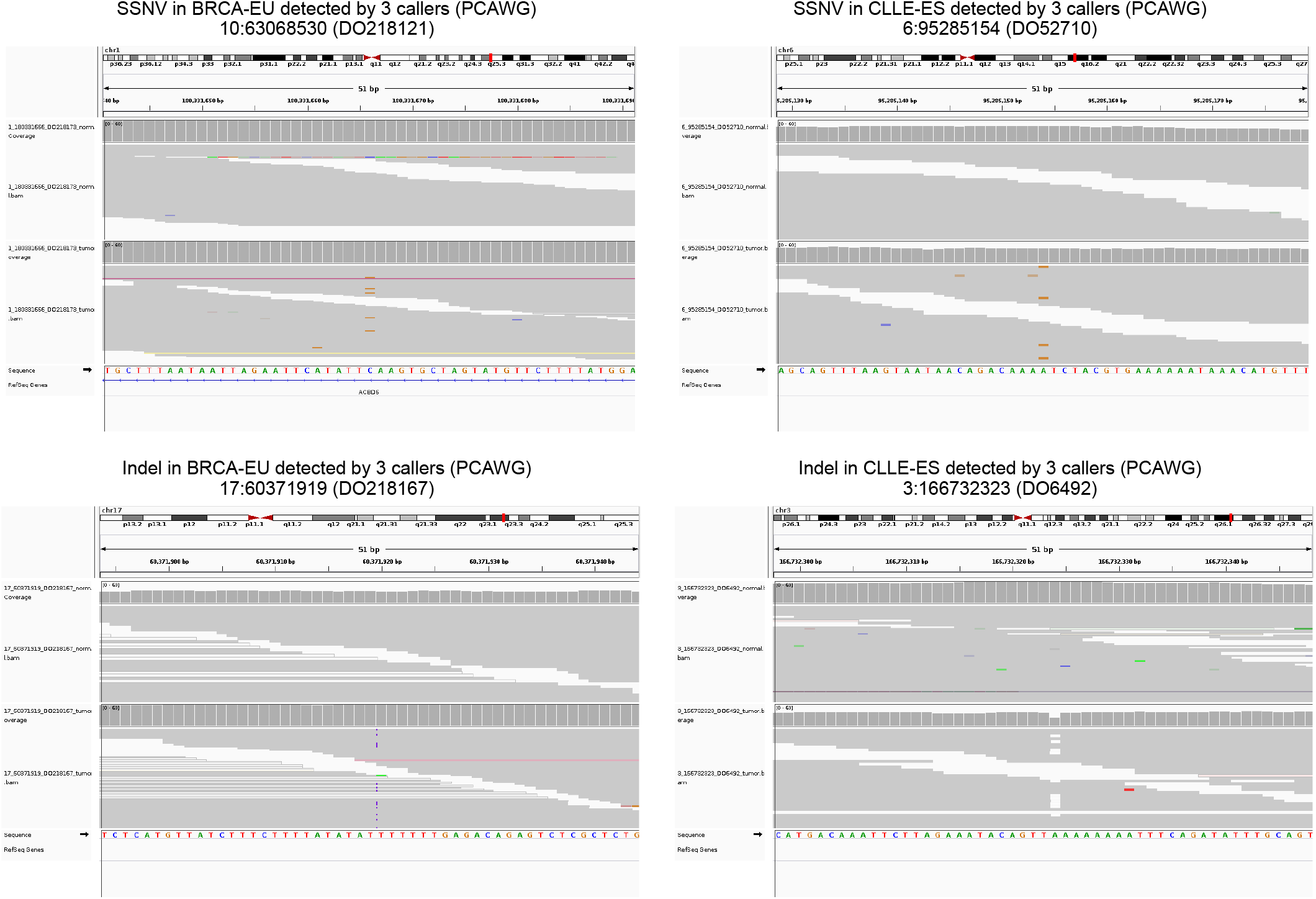
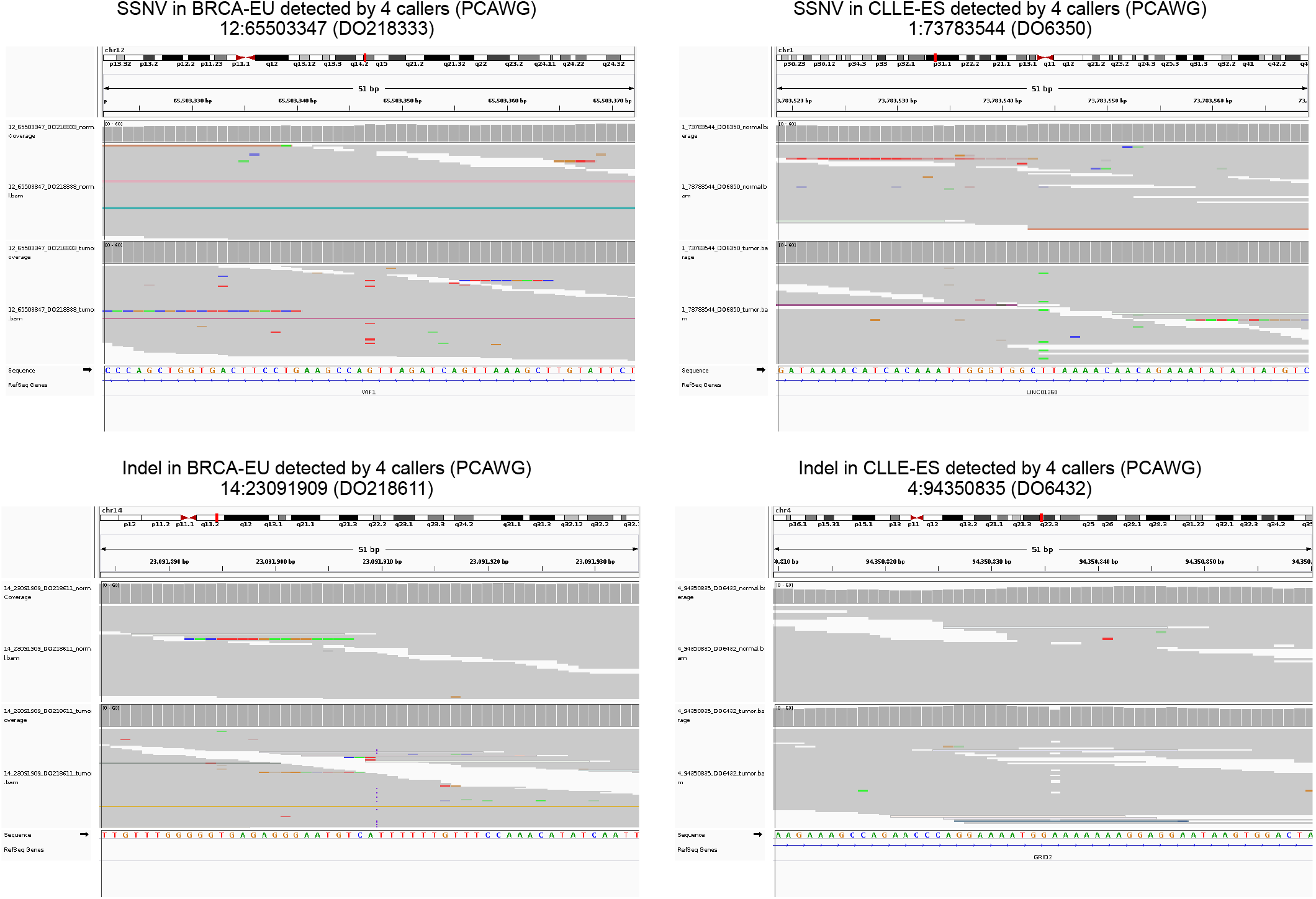
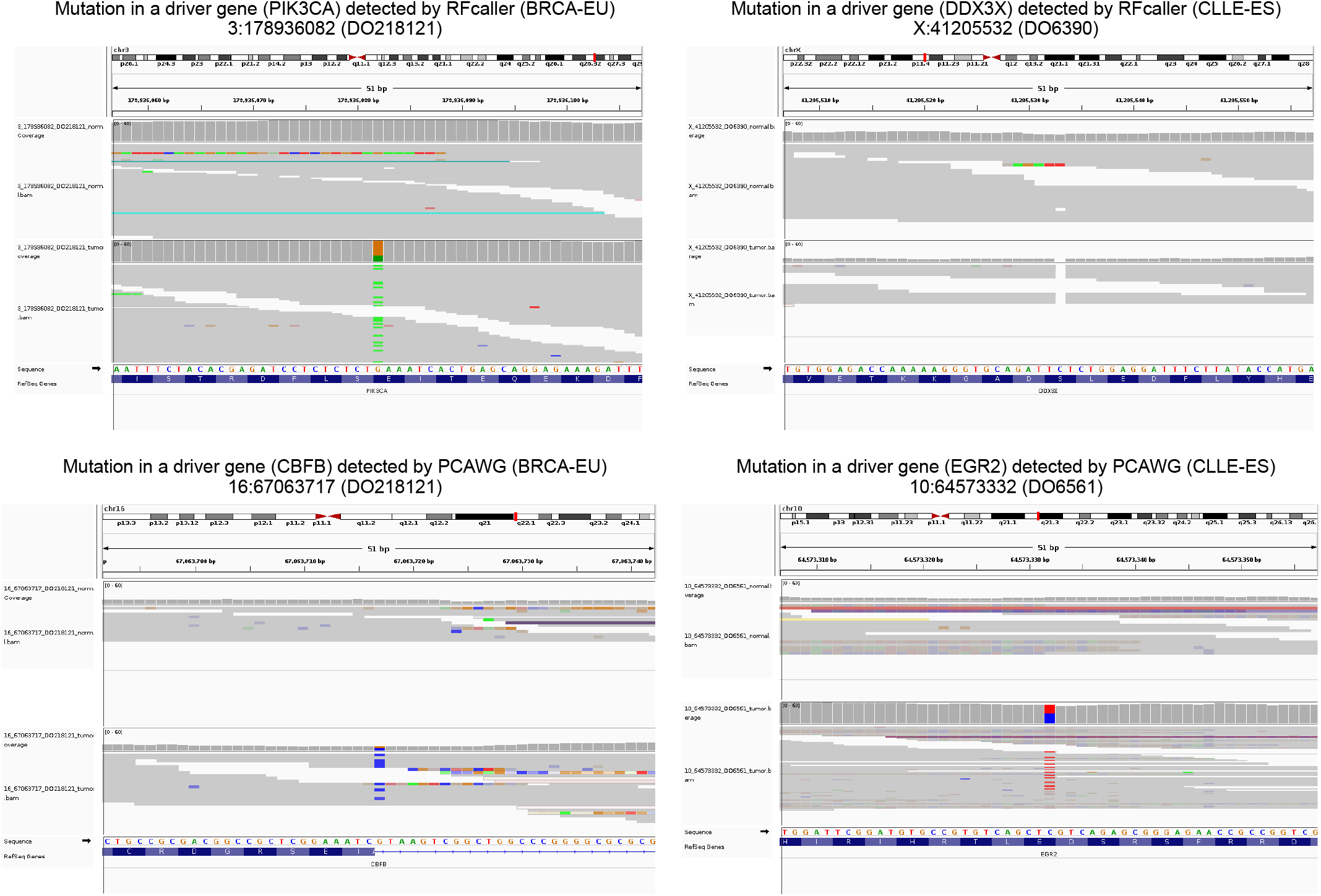
IGV screenshots corresponding to representative examples of mutations detected specifically by RFcaller o by the PCAWG pipelines.

**Supplementary Figure S4.**
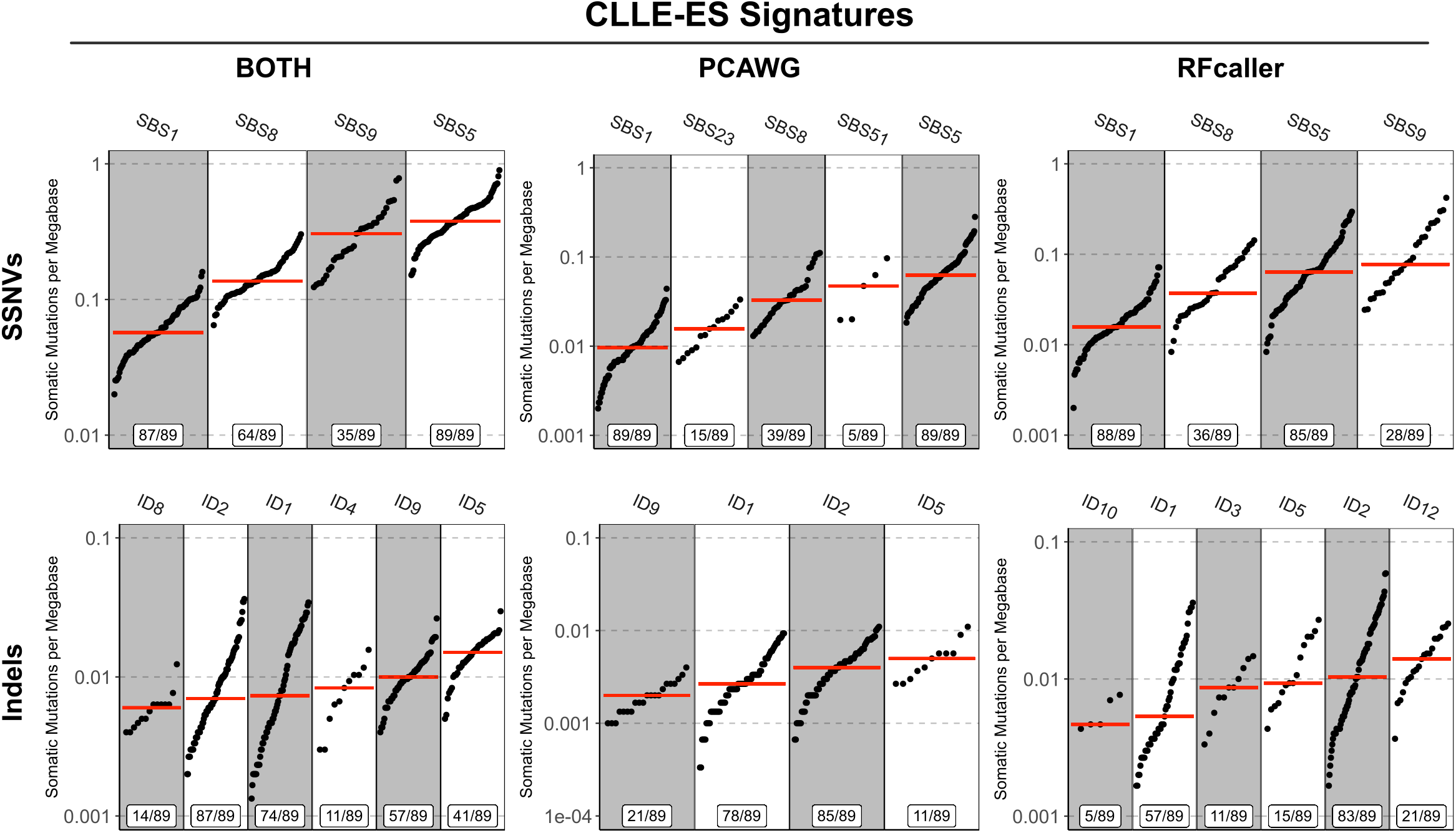
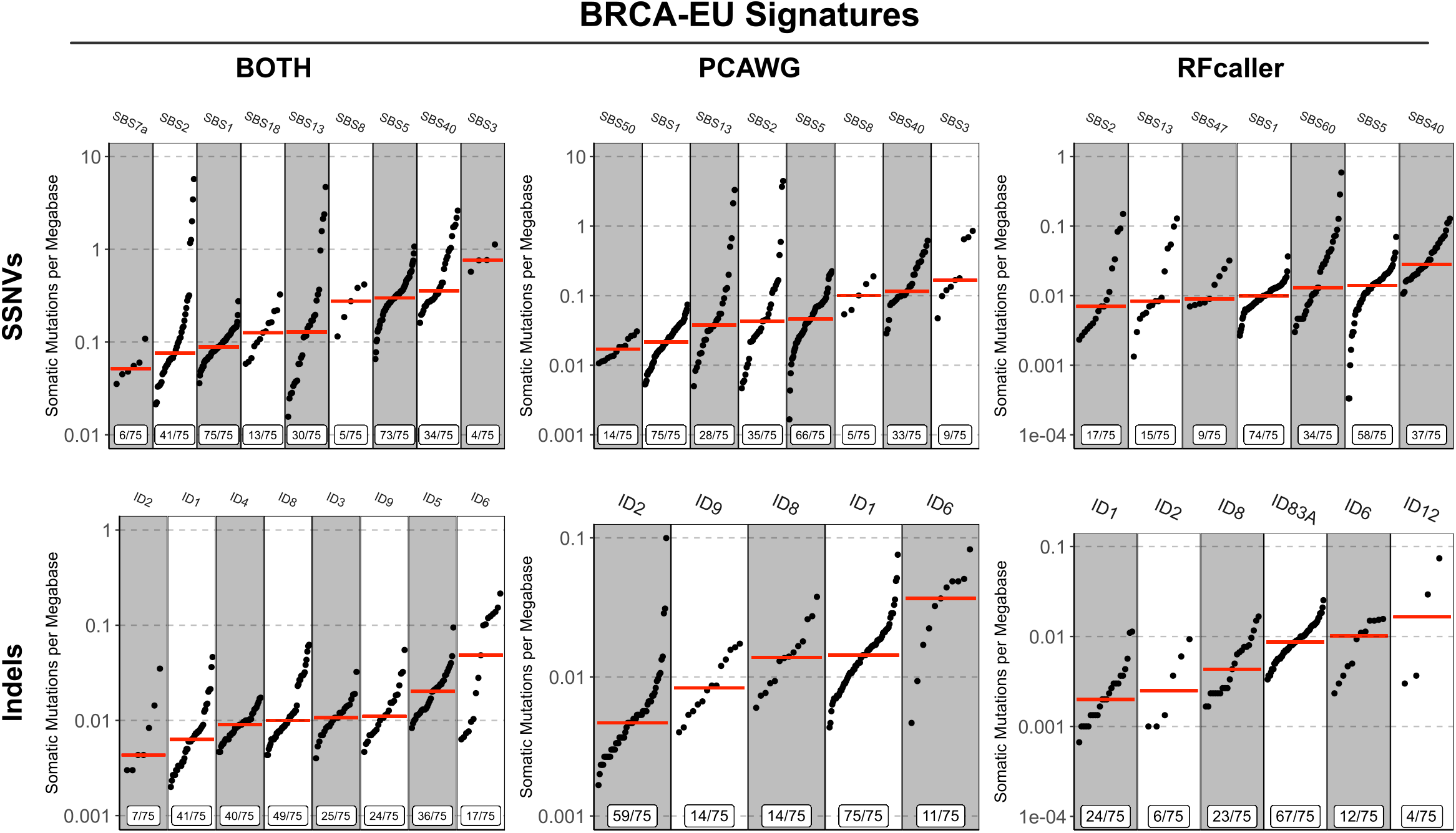
Mutational signatures extracted for CLLE-ES and BRCA-EU studies using the set of mutations detected by both pipelines and RFcaller and PCAWG-private mutations independently.

**Supplementary Figure S5.**
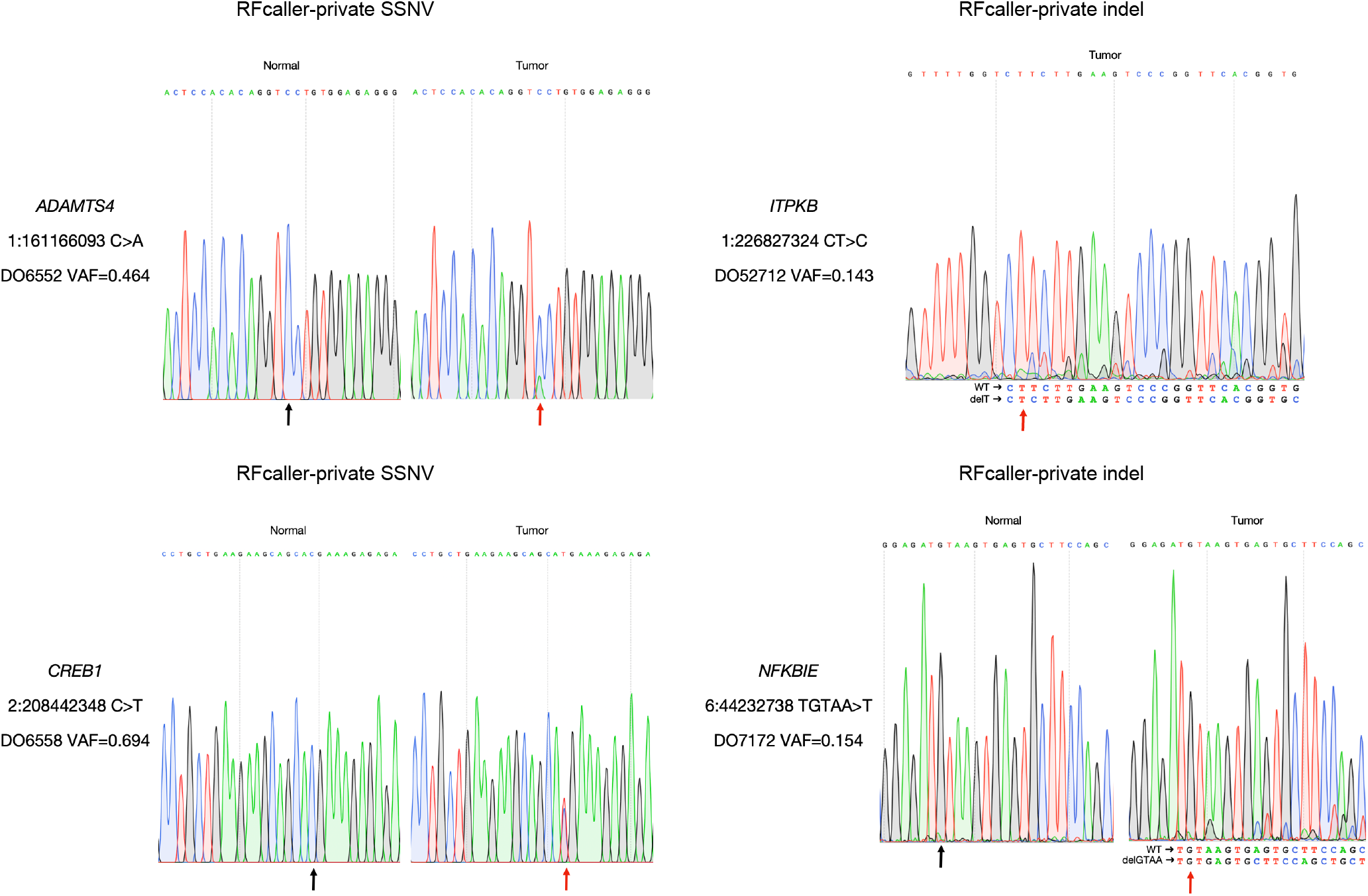
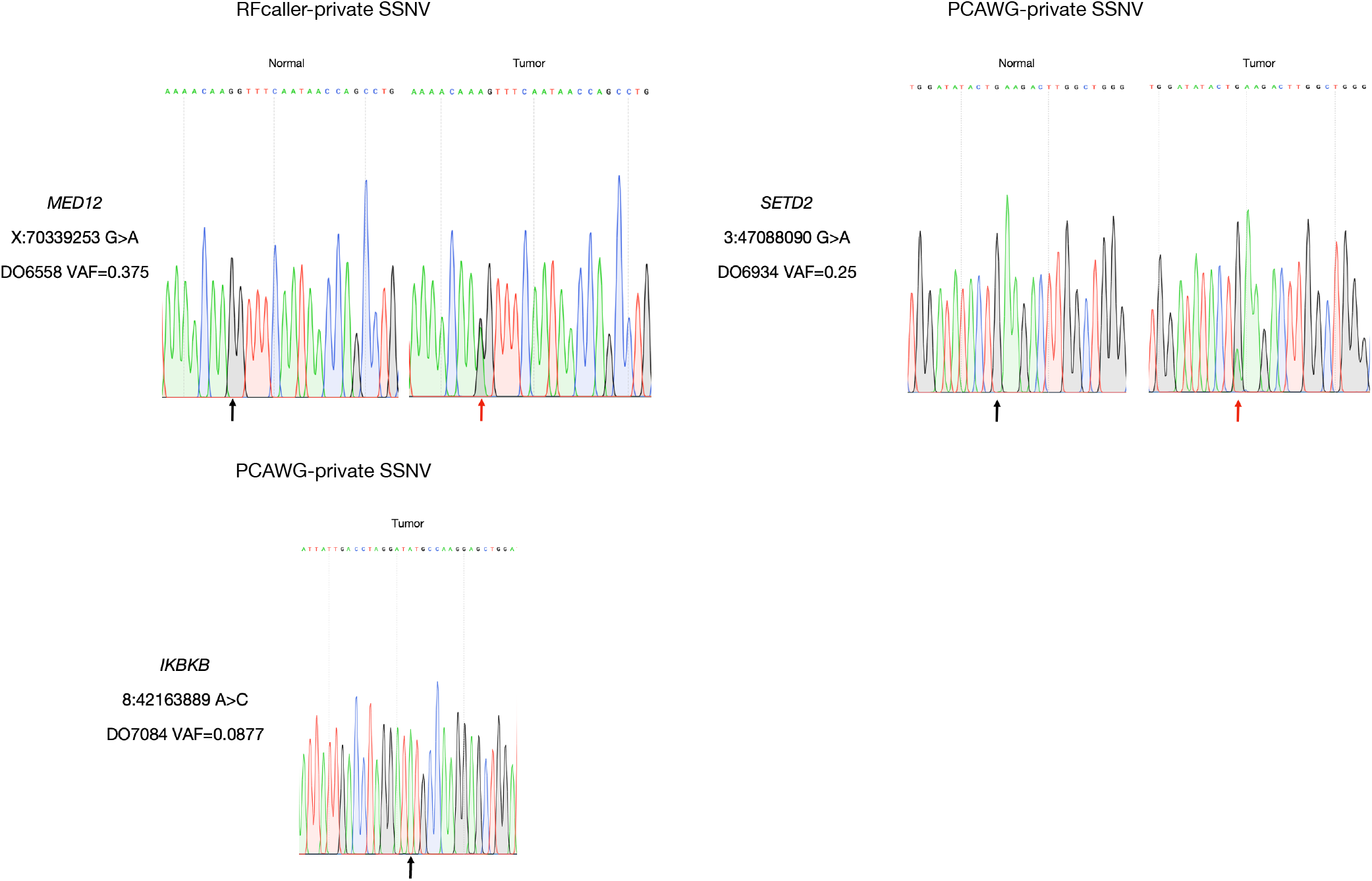
Electropherograms corresponding to Sanger verification of private mutations detected by RFcaller and PCAWG in CLL driver genes.

## Notes

https://github.com/xa-lab/RFcaller

https://hub.docker.com/repository/docker/labxa/rfcaller

